# The combined influence of viscoelasticity and adhesive cues on fibroblast spreading and focal adhesion formation

**DOI:** 10.1101/2021.02.17.430924

**Authors:** Erica Hui, Leandro Moretti, Thomas H. Barker, Steven R. Caliari

## Abstract

Tissue fibrosis is characterized by progressive extracellular matrix (ECM) stiffening and loss of viscoelasticity that ultimately results in reduced organ functionality. Cells bind to the ECM through integrins, where av integrin engagement in particular has been correlated with fibroblast activation into contractile myofibroblasts that drive fibrosis progression. There is a significant unmet need for *in vitro* hydrogel systems that deconstruct the complexity of native tissues to better understand the individual and combined effects of stiffness, viscoelasticity, and integrin engagement on fibroblast behavior. Here, we developed hyaluronic acid hydrogels with independently tunable cell-instructive properties (stiffness, viscoelasticity, ligand presentation) to address this challenge. Hydrogels with mechanics matching normal or fibrotic lung tissue were synthesized using a combination of covalent crosslinks and supramolecular interactions to tune viscoelasticity. Cell adhesion was mediated through incorporation of either RGD peptide or engineered fibronectin fragments promoting preferential integrin engagement via αvβ3 or α5β1. We showed that preferential αvβ3 engagement enabled human lung fibroblasts to assume a myofibroblast-like phenotype on fibrosis-mimicking stiff elastic hydrogels with increased spreading, actin stress fiber organization, and focal adhesion maturation as indicated by paxillin organization. In contrast, preferential α5β1 binding suppressed these metrics. Viscoelasticity, mimicking the mechanics of healthy tissue, largely curtailed fibroblast spreading and focal adhesion organization independent of adhesive ligand type, highlighting its role in preventing fibroblast activation. Together these results provide new insights into how mechanical and adhesive cues collectively guide disease-relevant cell behaviors.

## 1. Introduction

Tissue fibrosis is a pathological scarring process characterized by the excessive deposition of crosslinked extracellular matrix (ECM) proteins leading to progressive matrix stiffening and decreased viscoelasticity^25,35,48,61,79,82^. These aberrant changes in tissue mechanics detrimentally impact organ function, contributing to the role fibrosis plays in nearly half of all deaths in the developed world^33,52,81^. Reciprocal interactions between fibroblasts and their surrounding extracellular microenvironment actively drive a cascade of biochemical and biophysical signaling events to direct both normal and fibrogenic behaviors including adhesion, spreading, focal adhesion organization, and activation into fibrosis-promoting myofibroblasts^8,23,36,38,39,41^. However, delineating the specific environmental regulators of fibroblast behavior is difficult in multifaceted tissue milieus.

Numerous *in vitro* studies have used hydrogel biomaterials to deconstruct complex *in vivo* cellular microenvironments to better understand the individual and combined influence of biophysical factors such as stiffness and viscoelasticity on driving fibrogenic cell behaviors^4,11,12,14,16,18,27,83^. It is well understood that stiffer microenvironments can guide mechanotransduction by providing biophysical cues for fibroblast activation. Culturing cells atop substrates of increasing stiffness promotes increased spreading, actin stress fiber organization, and nuclear localization of transcriptional cofactors regulating the expression of fibrogenic genes encoding α-smooth muscle actin (α-SMA) and type I collagen^5,10,44,58,74,78,80,83^. While many studies of mechanotransduction use covalently-crosslinked hydrogels that behave as elastic solids, tissues are viscoelastic, meaning they exhibit both elastic solid and viscous liquid-like behaviors such as stress relaxation^25,40,85^. Seminal studies incorporating viscoelasticity into hydrogels showed that, compared to stiffness-matched elastic controls, cells displayed reduced spreading and expression of disease-relevant markers such as α-SMA with increasing loss modulus (viscoelasticity) due to reduced cellular contractility as a result of viscous dissipation^16,37^, highlighting the importance of viscoelasticity in disease mechanobiology.

While stiffness and viscoelasticity are well-established regulators of cell behavior, comparatively little attention has been paid to engineering hydrogels that can control cell adhesive interactions through specific integrin engagement. Integrins are transmembrane proteins composed of α and β subunits that bind to the ECM and serve as conduits for biochemical and mechanical signaling between cells and the ECM^42,67^. Importantly, integrin-based adhesions enable the conversion of complex biophysical cues, such as matrix mechanics and viscoelasticity, into chemical signals through mechanotransduction^3,19,34,62,72^. Integrin engagement and clustering facilitates the recruitment and formation of force-dependent focal adhesions (FAs) composed of proteins including paxillin, which play an important role in regulating cell behaviors such as spreading, contraction, migration, and differentiation^23,38,39,41,60,70^. As nascent cell-matrix adhesions (< 0.25 μm) mature into stable and larger FAs (1-5 μm), this strengthens integrin-FA-cytoskeletal linkages, facilitating actin polymerization and stress fiber organization, nuclear localization of transcriptional mechanoregulators, and the transcription of fibrogenic genes that ultimately results in dysregulated ECM production and organ failure^22,26,30,38,43,59,62^. While many synthetic hydrogels are engineered to support integrin-mediated cell attachment by incorporating the fibronectin-derived RGD peptide, this may inadvertently convolute mechanobiology studies due to its inefficient cell binding affinity compared to longer peptide or protein domains as well as its ability to non-specifically bind multiple integrin heterodimers^66^. Recent work has shown that provisional matrix proteins such as fibronectin (Fn) are upregulated during early stages of tissue remodeling and that integrin-specific Fn engagement (e.g., αvβ3 vs α5β1) caused by tension-stimulated conformational changes can influence fibrosis mechanoregulation^19,25,46,62^. In particular, engagement of the av integrin has been shown to promote integrin-mediated myofibroblast contractility^25,30,32,60^, mechanoactivation of latent transforming growth factor-beta 1 (TGF-βl)^2,24,34^, and expression and organization of a-SMA stress fibers, a hallmark of myofibroblast activation^5,20,68^.

While several studies, including from our group^37^, have highlighted the importance of stiffness and viscoelasticity in directing cell behavior, an approach to independently manipulate stiffness, viscoelasticity, and integrin engagement in a single system has not been developed. To address this challenge, we designed a phototunable viscoelastic hydrogel platform to deconstruct the complexity of native tissue toward understanding the individual and combined roles of cell-instructive cues including stiffness, viscoelasticity, and integrin-binding ligand presentation. We then used this system to determine how multiple mechanoregulatory cues work together to guide cellular behavior in the context of fibroblast activation.

## 2. Materials and Methods

### 2.1. NorHA synthesis

HA was functionalized with norbornene groups as previously described^28,37^. Sodium hyaluronate (Lifecore, 62 kDa) was converted to hyaluronic acid *tert-buty1* ammonium salt (HA-TBA) via proton exchange with Dowex 50W resin prior to being filtered, titrated to pH 7.05, frozen, and lyophilized. 5-norbornene-2-methylamine and benzotriazole-1-yloxytris-(dimethylamino)phosphonium hexafluorophosphate (BOP) were added dropwise to HA-TBA in dimethylsulfoxide (DMSO) and reacted for 2 hours at 25°C, quenched with cold water, dialyzed (molecular weight cutoff: 6-8 kDa) for 5 days, filtered, dialyzed for 5 more days, frozen, and lyophilized. The degree of modification was 31% as determined via proton nuclear magnetic resonance (^1^H NMR 500 MHz Varian Inova 500, **Figure S1**).

### 2.2. β-CD-HA synthesis

β-cyclodextrin modified hyaluronic acid (CD-HA) was synthesized by coupling synthesized 6-(6-aminohexyl)amino-6-deoxy-β-cyclodextrin (β-CD-HDA) to HA-TBA in anhydrous DMSO in the presence of BOP^37,65^. The amidation reaction was carried out at 25°C for 3 hours, quenched with cold water, dialyzed for 5 days, filtered, dialyzed for 5 more days, frozen, and lyophilized. The degree of modification was 28% as determined by ^1^H NMR (**Figure S2**).

### 2.3. Peptide synthesis

Thiolated adamantane peptide (Ad-KKK**C**G) was synthesized on Rink Amide MBHA high-loaded (0.78 mmol/g) resin using solid phase peptide synthesis as previously described^37^. The peptide was cleaved in 95% trifluoroacetic acid, 2.5% triisopropylsilane, and 2.5% H2O for 2-3 hours, precipitated in cold ether, dried, resuspended in water, frozen, and lyophilized. Synthesis was confirmed via matrix-assisted laser desorption/ionization (MALDI) mass spectrometry (**Figure S3**).

### 2.4. Recombinant fibronectin fragments

Recombinant fibronectin fragments of the ninth and tenth type III repeat units (FnIII_9_ and FnIII_10_) were designed to preferentially bind a5β1 or avβ3 integrin heterodimers as previously described^25,47,53^. Briefly, to promote a5β1 binding, FnIII_9_ was thermodynamically stabilized through a leucine to proline point mutation at position 1408, which has demonstrated stabilization of the spatial orientation of the RGD motif on FnIII_10_ and the synergy site PHSRN on the FnIII_9_, increasing selectivity to β1 integrins^15^. While this fragment still supports avβ3 binding, it has greater α5β1 integrin-binding affinity (K_D_ ~ 12 nM for α5β1 versus ~ 40 nM for αvβ3)^15^. We have referred to this fragment as ‘Fn9*10’ throughout the manuscript. For avβ3 integrin binding specificity, four glycine residues were inserted into the liner region between FnIII_9_ and FnIII_10_ to disrupt a5β1 binding by increasing the separation between the RGD and PHSRN sites. This fragment is denoted ‘Fn4G’. Both fibronectin fragments contained N-terminal cysteine residues to enable thiol-ene coupling to the HA hydrogels.

### 2.5. HA hydrogel fabrication

Thin film hydrogels (18 x 18 mm, ~ 100 μm thickness) were fabricated on thiolated coverslips via ultraviolet (UV)-light mediated thiol-ene addition, similar to previously established methods^37^. ‘Soft’ and ‘stiff’ hydrogel formulations were designed to match normal (Young’s modulus or stiffness ~ 1 kPa) and fibrotic (~ 15 kPa) stiffnesses respectively^7,25,48,79^. Covalently-crosslinked soft (2 wt% NorHA) and stiff (6 wt% NorHA) elastic hydrogels formulations were crosslinked with dithiothreitol (DTT, thiol-norbornene ratios of 0.22 and 0.35 for soft and stiff groups, respectively). Soft (2 wt% NorHA-CDHA) and stiff (6 wt% NorHA-CDHA) viscoelastic hydrogels were fabricated through a combination of covalent and physical crosslinking. NorHA and DTT (covalent crosslinks, thiol-norbornene ratios of 0.35 and 0.55 for soft and stiff groups, respectively) were combined with CD-HA and thiolated adamantane (Ad) peptides (supramolecular guest-host interactions between CD and Ad, 1:1 molar ratio of CD to Ad). Cell adhesion was enabled in all hydrogel groups through incorporation of either 1 mM RGD peptide (GCGYGRGDSPG, Genscript) or 2 μM thiolated Fn fragments (Fn9*10 or Fn4G). Hydrogel solutions were photopolymerized (365 nm, 5 mW/cm^2^) between coverslips in the presence of 1 mM lithium acylphosphinate (LAP) photoinitiator for 2 minutes and swelled in PBS overnight at 37°C before subsequent experiments.

### 2.6. Mechanical characterization

Hydrogel rheological properties were tested on an Anton Paar MCR 302 rheometer using a cone-plate geometry (25 mm diameter, 0.5°, 25 μm gap). *In situ* gelation via 2 minute UV light irradiation (5 mW/cm^2^) was tracked using oscillatory time sweeps (1 Hz, 1% strain) followed by oscillatory frequency sweeps (0.001-10 Hz, 1% strain) and cyclic stress relaxation and recovery tests alternating between 0.1% and 5% strain. Nanoindentation tests were performed using Optics11 Piuma and Chiaro nanoindenters on hydrogels swollen in PBS for at least 24 hours to determine hydrogel mechanical characteristics. A 25 μm diameter spherical borosilicate glass probe attached to a cantilever with a spring constant of 0.5 N/m was used during testing. For each indentation, the loading portion of the generated force versus distance indentation curves was used to determine the Young’s modulus by applying the Hertzian contact mechanics model and assuming a Poisson’s ratio of 0.5. Topography was mapped through matrix indentations. The Optics11 nanoindenter software also features a dynamic operational mode to enable dynamic mechanical analysis (DMA)-like measurements through mechanical oscillations. DMA measurements were performed to study timedependent viscoelasticity (G′ and G″) of swollen hydrogels via frequency sweeps (0.1-10 Hz) and time-dependent force relaxation tests.

### 2.7. Cell culture

Human lung fibroblasts (hTERT T1015, abmgood) were used between passages 7-12 and culture medium was changed every 2-3 days (Gibco Dulbecco’s Modified Eagle Medium (DMEM) supplemented with 10 v/v% fetal bovine serum (FBS) and 1 v/v% antibiotic antimycotic (1,000 U/mL penicillin, 1,000 μg/mL streptomycin, and 0.25 μg/mL amphotericin B)). Normal human lung fibroblasts (CC-2512, Lonza) were used between passages 3-5 for paxillin experiments and culture medium was changed every 2-3 days (Lonza FBM Basal Medium supplemented with 2 v/v% fetal bovine serum (FBS), 0.1 v/v% human recombinant insulin (1-20 μg/mL), 0.1 v/v% recombinant human fibroblast growth factor-B (rhFGF-B, 0.5-5 ng/mL), and 0.1 v/v% gentamicin sulfate amphotericin B (GA-1000, 30 μg/mL gentamicin and 15 ng/mL amphotericin)). Swelled hydrogels were sterilized in untreated 6-well plates via germicidal UV irradiation for at least 2 hours and incubated in culture medium for at least 30 minutes prior to cell seeding. Cells were seeded at 2 x 10^4^ cells/hydrogel (18 x 18 mm).

### 2.8. Immunostaining, imaging, and analysis

Cell-seeded hydrogels were fixed in 10% neutral-buffered formalin for 15 minutes, permeabilized in PBST (0.1% Triton X-100 in PBS) for 10 minutes, and blocked in 3% bovine serum albumin (BSA) in PBS for at least 1 hour at 25°C. To visualize focal adhesions (FAs), cells were fixed using a microtubule stabilization buffer for 10 minutes at 37°C before blocking. Hydrogels were then incubated overnight at 4°C with primary antibodies. Primary antibodies used in this work included paxillin (mouse monoclonal anti-paxillin B-2, Santa Cruz Biotechnology, sc-365379, 1:500) to visualize FA formation and α-smooth muscle actin (α-SMA, mouse monoclonal anti-α-SMA clone 1A4, Sigma-Aldrich, A2547, 1:400). Hydrogels were washed three times using PBS and incubated with secondary antibodies (AlexaFluor 488 goat anti-rabbit IgG or AlexaFluor 555 goat anti-mouse IgG, Invitrogen, 1:600-800) and/or rhodamine phalloidin (Invitrogen, R415 1:600) to visualize Factin for 2 hours in the dark at 25°C. Hydrogels were rinsed three times with PBS and incubated with a DAPI nuclear stain (Invitrogen, D1306, 1:10000) for 1 minute before washing with PBS. Images were taken on a Zeiss AxioObserver 7 inverted microscope. Cell spread area and cell shape index were determined using a CellProfiler (Broad Institute, Harvard/MIT) pipeline modified to include adaptive thresholding. Cell shape index determines the circularity of the cell, where a line and a circle have values of 0 and 1, respectively, and was calculated using the formula:

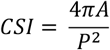

where *A* is the cell area and *P* is the cell perimeter. For FA analysis, cells stained with paxillin were imaged using a 40x oil objective. FA count, area, and fluorescence intensity were quantified via the Focal Adhesion Analysis Server (FAAS)^6^ automated imaging processing pipeline using a 4.5 threshold and minimum pixel size of 25.

### 2.9. Statistical analysis

For mechanical characterization, at least 3 technical replicates were performed and the data are presented as mean ± standard deviation. For statistical comparisons between hydrogel groups, two-way ANOVA with Tukey’s HSD post hoc analysis (more than two experimental groups) were performed. All experiments included at least 3 replicate hydrogels per experimental group. Box plots of single cell data include median/mean indicators as well as error bars corresponding to the lower value of either the 1.5*interquartile range or the maximum/minimum value, with data points outside the 1.5*interquartile range shown as open circles. Statistically-significant differences are indicated by *, **, or *** corresponding to *P* < 0.05, 0.01, or 0.001 respectively.

## 3. Results and Discussion

### 3.1. Hydrogels were designed to independently control stiffness, viscoelasticity, and presentation of integrin-binding adhesive sites

Hyaluronic acid (HA) hydrogels representing normal (G′ ~ 0.5 kPa) and fibrotic (G′ ~ 5 kPa) lung tissue mechanics were fabricated with a combination of covalent crosslinks and supramolecular guest-host interactions to impart viscous properties (**Figure 1**)^37,64,65^. HA was chosen as the hydrogel backbone for its ability to be chemically modified with various functional groups to achieve a range of viscoelastic properties covering healthy and diseased soft tissue, as shown in previous work by our lab and others^9,13,28,37,45,69^. Stiffness was controlled primarily through adjusting the concentration of HA and the ratio of dithiol crosslinker to norbornene groups on HA. Several methods to incorporate viscoelasticity into material systems have been developed, including the addition of sterically entrapped high molecular weight linear polymers to introduce viscosity^1,16^, covalent adaptable networks^51,55,73^, physical crosslinking of natural polymers (e.g., alginate^17,84^, collagen^56,57^) for modulation of stress relaxation properties, and supramolecular crosslinking chemistries (e.g., host-guest complexes^37,63,65,77^). In this work, the addition of supramolecular guest-host interactions between β-cyclodextrin HA (CD-HA) and thiolated adamantane (Ad) peptides (1:1 molar ratio of CD to Ad), where the hydrophobic Ad guest moiety has a high affinity for the hydrophobic interior of CD, introduced viscous characteristics into the system^37,65^. Elastic hydrogel substrates contained only covalent crosslinks, while viscoelastic substrates included a combination of covalent and supramolecular interactions.

**Figure 1.**
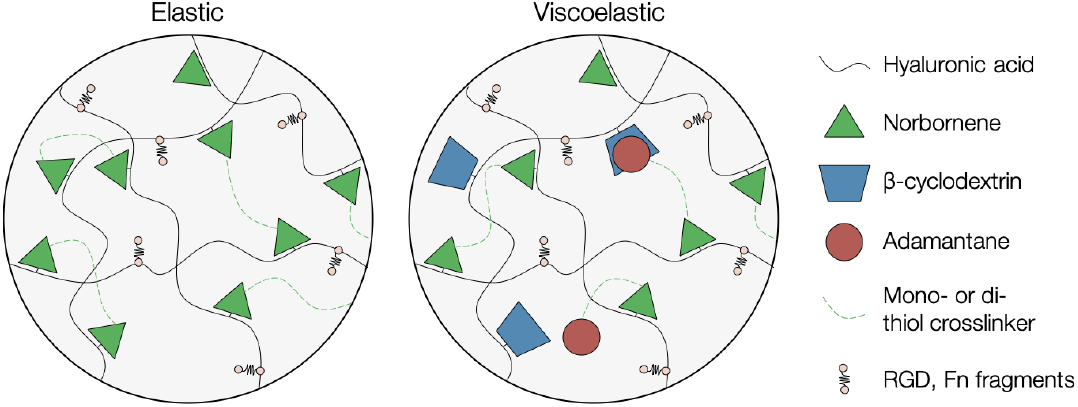
Schematic of elastic and viscoelastic hyaluronic acid hydrogel design. Covalent crosslinks between norbornenes and di-thiol crosslinkers are formed via light-mediated thiol-ene addition to create elastic hydrogel networks. A combination of covalent crosslinking and supramolecular guest-host interactions between cyclodextrins and thiolated adamantane groups confer viscous characteristics to the viscoelastic system. Thiolated adhesive ligands (RGD or Fn fragments) were also incorporated during hydrogel formation.

While HA is a natural ECM component and interacts with cell surface receptors including CD44 and RHAMM in its unmodified forms, it does not support integrin binding, allowing customization of these interactions in our hydrogel design^11,21^. In addition to controlling hydrogel stiffness and viscoelasticity by modulating crosslinking as described above, we hypothesized that we could also dictate cellular adhesion through the incorporation of either thiolated RGD peptide or Fn fragments designed to preferentially bind αvβ3 (Fn4G) or α5β1 (Fn9*10) integrins^25,47,53^. Preferential α5β1 engagement in Fn9*10 is engineered by stabilizing the spatial proximity of the PHSRN synergy site on Fnlll9 with the RGD on FnIII_10_, although Fn9*10 can also bind αvβ3^15^. Insertion of a four glycine spacer between FnIII_9_ and FnIII_10_ in Fn4G abrogates simultaneous binding to both the PHSRN and RGD sequences necessary for α5β1 engagement, leading to preferential αvβ3 binding^15^. Since the RGD peptide does not contain the PHSRN synergy sequence, we anticipate that it would also preferentially engage αvβ3 over α5β1. Overall, the modular hydrogel design allows independent control of HA content, crosslinking type and density, and adhesive ligand incorporation to enable simultaneous tuning of stiffness, viscoelasticity, and integrin engagement.

### 3.2. Incorporation of fibronectin-related adhesive ligands did not impact hydrogel mechanics

We next wanted to determine if incorporating different adhesive ligands would impact the ability to independently control hydrogel stiffness and viscoelasticity. Hydrogel mechanics were examined through *in situ* oscillatory shear rheology (**Figure 2A,B**) and nanoindentation of PBS-swollen hydrogels (**Figure 2C,D**). Rapid *in situ* gelation kinetics for all hydrogel experimental groups was confirmed via rheology (**Figure S4**). The introduction of fibronectin-based adhesive ligands did not affect overall mechanics; similar storage and loss moduli were observed for all groups compared to RGD-containing hydrogels. Target mechanical values for ‘soft’ and ‘stiff’ groups corresponding to normal (elastic modulus, E ~ 1 kPa) and fibrotic (E ~ 15 kPa) lung tissue were successfully reached. As expected, the viscoelastic hydrogel design led to increased viscous properties as evidenced by higher loss moduli (G″) that were within an order of magnitude of the storage moduli (G′), analogous to normal soft tissue like lung and liver^61^.

**Figure 2.**
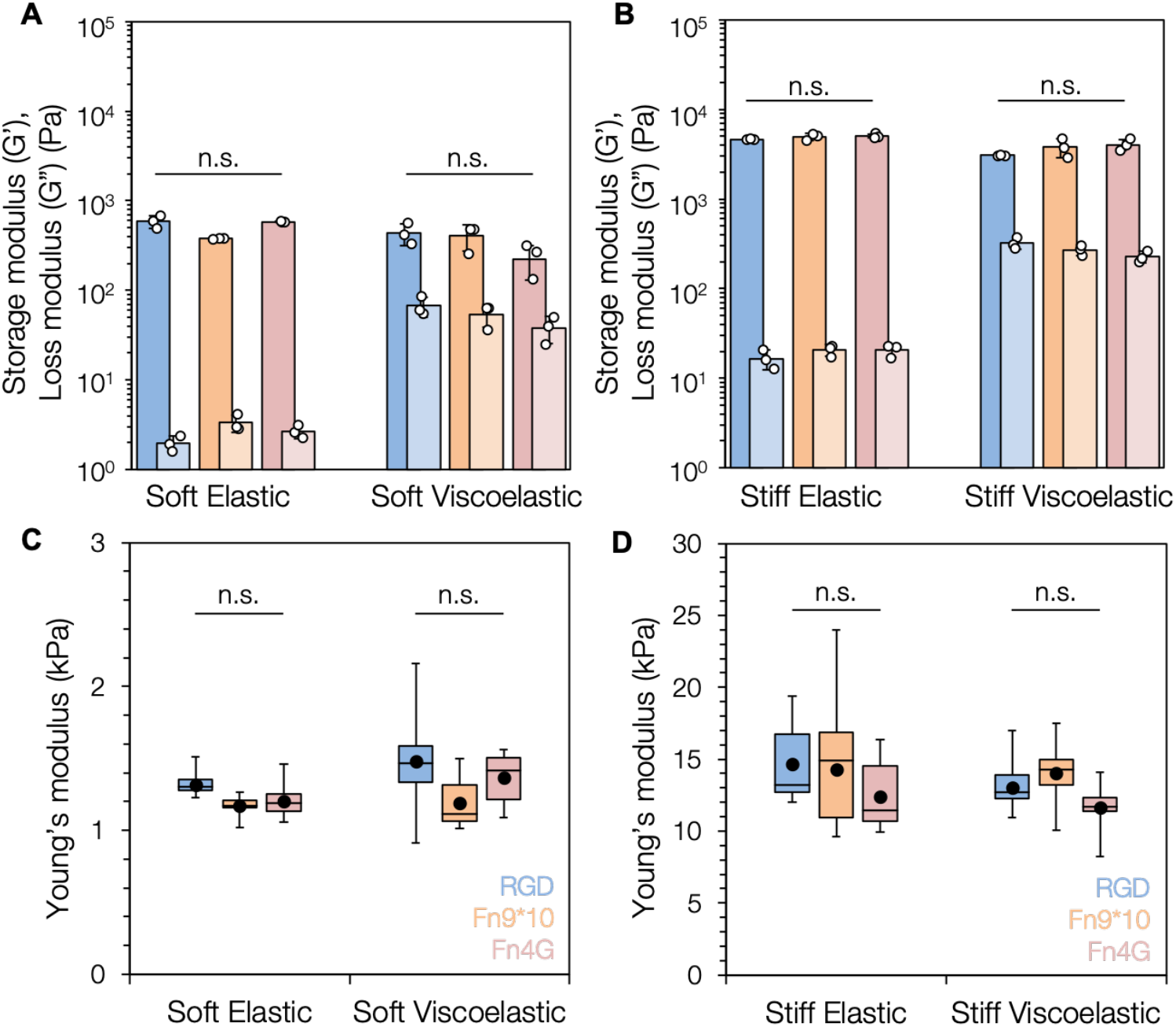
Mechanical characterization of viscoelastic hydrogels. (A) Average values of soft elastic and soft viscoelastic storage (G′) and loss (G″) moduli measured at a constant frequency (1 Hz) and strain (1%), characterized by oscillatory shear rheology, show clear differences in loss moduli between elastic and viscoelastic groups but no significant differences as a function of adhesive ligand type. (B) Average values of stiff elastic and stiff viscoelastic storage (G′) and loss (G″) moduli measured at a constant frequency (1 Hz) and strain (1%), characterized by oscillatory shear rheology, show similar trends to the soft hydrogel groups. (C) Box and whisker plots of soft elastic and soft viscoelastic Young’s moduli (E) of swollen hydrogels, characterized via nanoindentation, demonstrate equivalent Young’s moduli (stiffnesses) for all groups. (D) Box and whisker plots of stiff elastic and stiff viscoelastic Young’s moduli (E) of swollen hydrogels, characterized via nanoindentation, show similar trends to the soft hydrogel groups. Box plots of indentation data show median *(line),* mean *(filled black circle),* and have error bars corresponding to the lower value of either 1.5*interquartile range or the maximum/minimum value, with individual data points outside the 1.5*interquartile range shown as open circles. At least 3 hydrogels were tested per experimental group.

Viscoelastic substrates also displayed tissue-relevant frequencydependent mechanical responses as measured by both rheology (**Figures 3, S5**) and DMA-like nanoindentation measurements (**Figure S6**); at lower frequencies (longer time scales), the ability for guest-host interactions to re-organize and re-associate resulted in more solid-like behavior, whereas at higher frequencies (shorter time scales) guest-host interactions were disrupted with less time for complex reformation^37,65^. Stress relaxation, a key feature of viscoelastic materials, was demonstrated by observation of time-dependent decreases in storage modulus only in viscoelastic substrates when a constant strain (5%) was applied (**Figure S7**). The frequency-dependent relaxation behavior observed for the viscoelastic groups relates to cell-relevant time scales; cells are able to respond to force oscillations and exert traction forces on the order of seconds to minutes at a frequency of around 0.1-1 Hz^14,16,17^. Elastic hydrogels consisting of only stable covalent crosslinks did not display stress relaxation over time.

**Figure 3.**
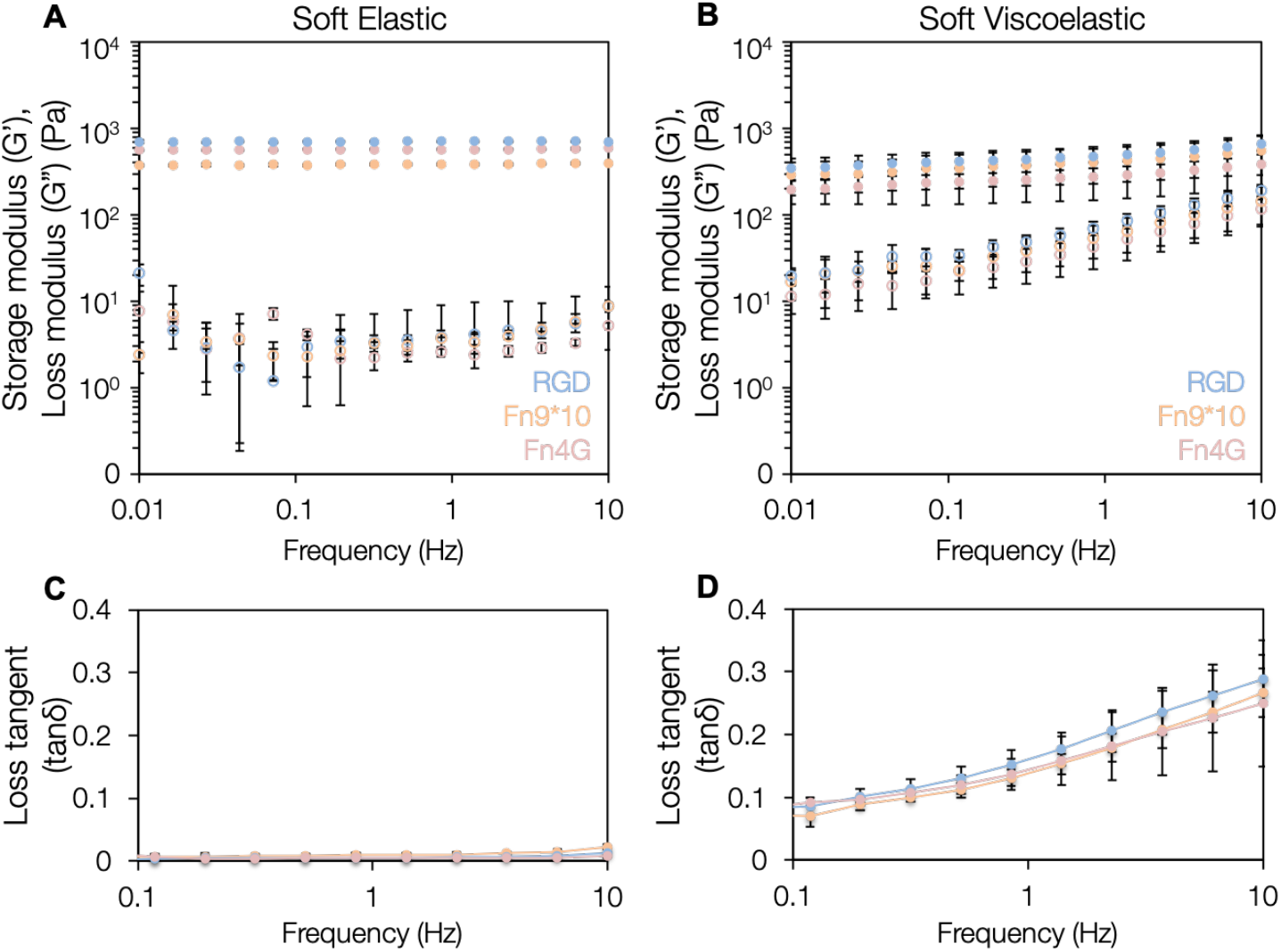
Frequency-dependent behavior of viscoelastic hydrogels. (A) Elastic hydrogels showed frequency-independent behavior, with storage (G′, *closed circles)* and loss moduli (G″, *open circles)* remaining relatively constant. (B) In contrast, viscoelastic hydrogels displayed frequency-dependent behavior with increasing loss moduli *(open circles)* at increasing frequencies. (C) Loss tangent (tanδ) values, which represent the ratio of viscous to elastic mechanical properties (G″/G′), remained relatively constant and close to 0 for all elastic hydrogels. (D) In contrast, loss tangent values were elevated for viscoelastic groups across all frequencies tested and increased at higher frequencies. Similar trends were seen for the stiff groups (Figure S5). Similar results for swollen hydrogel samples were measured using dynamic mechanical analysis (DMA)-like nanoindentation (Figure S6). 3 hydrogels were tested per experimental group.

### 3.3. Fibroblast spreading is influenced by both viscoelasticity and adhesive ligand type

After validating that hydrogels incorporating different adhesive ligands could be synthesized in both elastic and viscoelastic forms with overall stiffness matching normal and fibrotic tissue, we sought to confirm that our hydrogel formulations would support equivalent cell adhesion. We quantified the number of fibroblasts attached to the hydrogels after one day and confirmed that all formulations supported similar levels of adhesion (**Figure S8**). Notably, hydrogels containing only 2 μM Fn fragments allowed equivalent fibroblast attachment to hydrogels with 1 mM RGD peptide. Previous work using these fragments has also shown robust cell attachment using concentrations of this magnitude^15,47^. In contrast, short linear RGD peptides have previously been shown to be around 1000 times less effective in cell attachment compared to fibronectin^31^.

We next investigated the combined influence of stiffness, viscoelasticity, and adhesive ligand presentation on fibroblast spread area and shape. We used increased spreading as a proxy for increased cell contractility and myofibroblast activation as previously observed in many *in vitro* systems^5,16,29,54,83^. Human lung fibroblasts were seeded atop hydrogels and cultured for three days. We then quantified fibroblast spread area and cell shape index, a measure of cell circularity between 0 and 1 where 0 is a line and 1 is a circle (**Figure 4**). For the RGD-presenting hydrogels, cells showed greater spreading (2590 ± 670 μm^2^) on stiff elastic groups compared to smaller morphologies on soft (1210 ± 650 μm^2^) and stiff (1110 ± 510 μm^2^) viscoelastic groups, similar to results observed in previous studies^37^. The promotion of α5β1 engagement largely blunted the stiffness-dependent spreading response with fibroblasts showing reduced spreading and more rounded morphologies across all hydrogel groups regardless of stiffness or viscoelasticity (average spread area on Fn9*10 hydrogels: 780 ± 490 μm^2^), similar to previous findings with alveolar epithelial cell spreading^8,50^. Hydrogels supporting αvβ3-specific integrin engagement promoted similar levels of spreading to RGD-modified substrates, although increased spreading was observed even on soft elastic substrates (2060 ± 640 μm^2^). However, cells displayed decreased spreading and remained rounded on viscoelastic hydrogels regardless of stiffness. Additionally, we used nanoindentation to measure apical fibroblast stiffness on the different hydrogel formulations and found that fibroblasts were significantly stiffer on stiff elastic hydrogels where they preferentially engaged αvβ3 (RGD, Fn4G groups), but not on Fn9*10-modified hydrogels (**Figure S9**).

**Figure 4.**
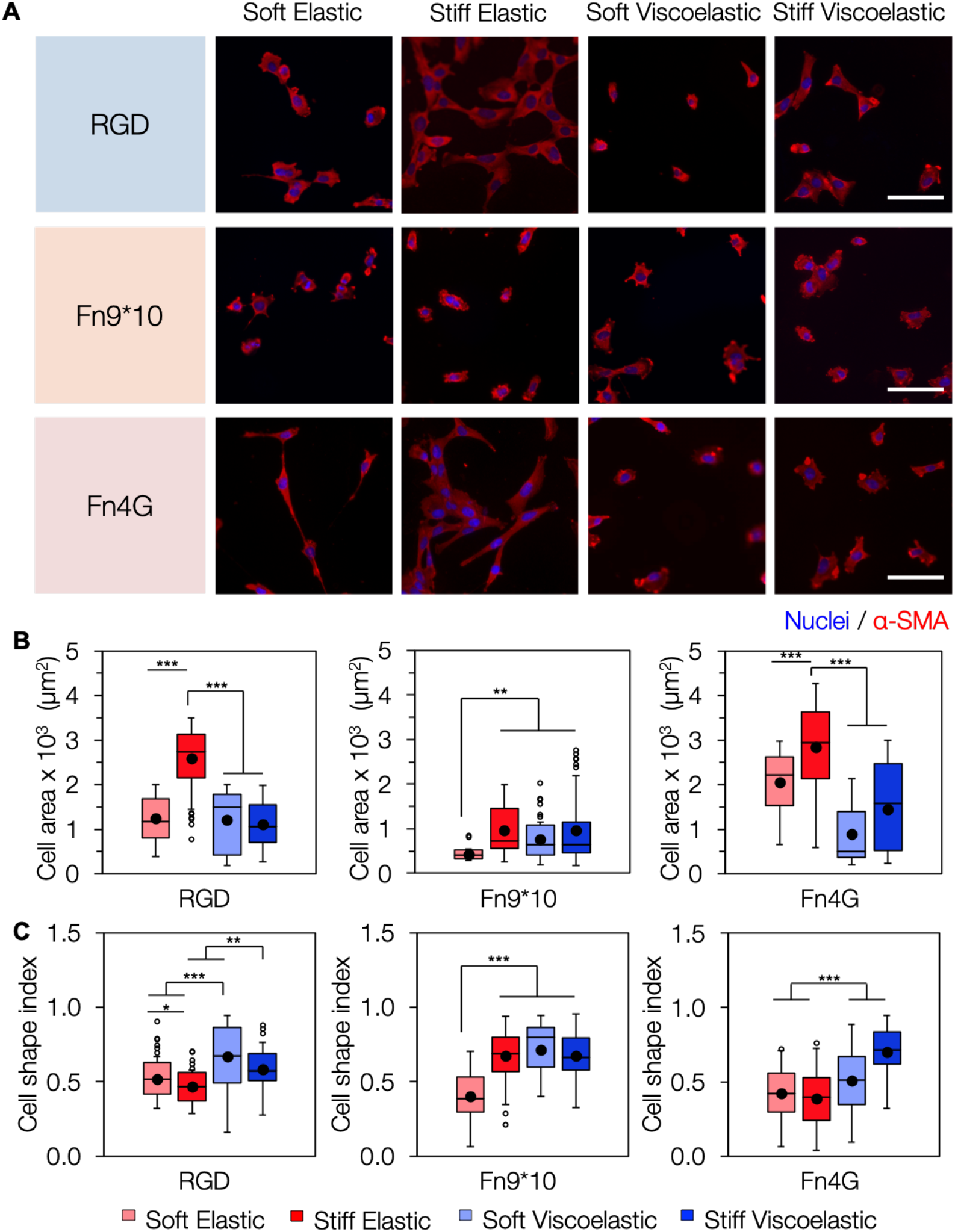
Fibroblast spreading is influenced by both viscoelastic mechanics and adhesive ligand type. (A) Human lung fibroblasts were cultured for 3 days on soft or stiff elastic and viscoelastic hydrogel groups modified with either RGD or fibronectin fragments preferentially engaging α5β1 or αvβ3. B) Fibroblasts preferentially binding αvβ3 (RGD, Fn4G) displayed increased spread area on elastic groups regardless of stiffness, but viscoelasticity suppressed spreading on all groups. C) Cell shape index showed correlative results with spreading as smaller fibroblasts remained elongated (lower cell shape index) while larger fibroblasts assumed a more spread, activated morphology. Box plots of single cell data show median *(line),* mean *(filled black circle),* and have error bars corresponding to the lower value of either 1.5*interquartile range or the maximum/minimum value, with data points outside the 1.5*interquartile range shown as open circles. *Scale bars:* 100 μm, *: *P* < 0.05, **: *P* < 0.01, ***: *P* < 0.001. 3 hydrogels were tested per experimental group (50-600 cells total).

### 3.4. Preferential αvβ3 integrin engagement promotes actin stress fiber organization and larger focal adhesion formation

The differences in fibroblast spreading observed as a function of stiffness, viscoelasticity, and adhesive ligand motivated us to more completely understand potential differences in cytoskeletal organization, particularly actin stress fiber formation and focal adhesion maturation. First, we qualitatively evaluated the level of actin stress fiber organization as well as the organization of paxillin, a prominent focal adhesion (FA) adaptor protein that has been implicated in regulating cytoskeletal organization^49,71,75,76^, in fibroblasts seeded on hydrogels (**Figure 5**). We found that actin organization was strongly correlated to spread area, with fibroblasts on stiff elastic Fn4G hydrogels engaging primarily αvβ3 mostly displaying actin organized into stress fibers. In contrast, fibroblasts on Fn9*10 hydrogels showed few organized stress fibers, even on stiff elastic hydrogels mimicking fibrotic tissue. Notably, actin stress fiber organization was absent in the vast majority of fibroblasts cultured on soft or stiff viscoelastic hydrogels regardless of adhesive ligand functionalization.

**Figure 5.**
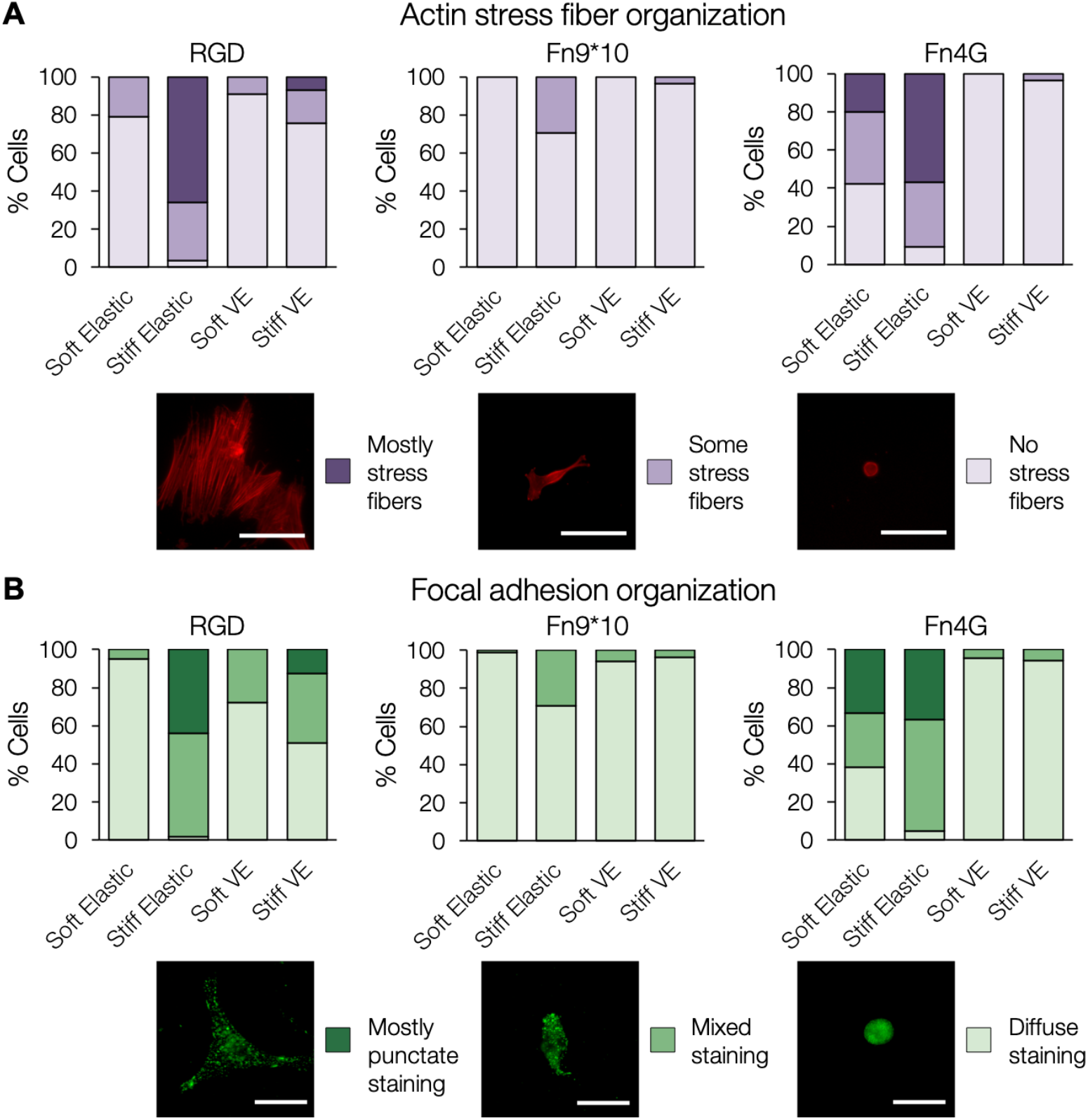
Qualitative analysis of actin stress fiber and focal adhesion organization. (A) Percentage of human lung fibroblasts showing various levels of F-actin stress fiber organization as indicated by the representative images. More actin stress fibers were observed in fibroblasts on stiff elastic hydrogels, especially for groups preferentially binding αvβ3 (RGD, Fn4G) while viscoelasticity suppressed stress fiber formation across all ligand groups. *Scale bars:* 100 μm. 3 hydrogels were tested per experimental group (60-110 cells total). (B) Percentage of human lung fibroblasts showing various levels of paxillin organization as indicated by the representative images. Similarly to the results in (A), more punctate paxillin staining was observed in fibroblasts on stiff elastic hydrogels, especially for groups preferentially binding αvβ3 (RGD, Fn4G) while viscoelasticity suppressed focal adhesion maturation across all ligand groups. *Scale bars:* 50 μm. 3 hydrogels were tested per experimental group (40-130 cells).

On RGD-containing hydrogels punctate focal adhesion organization, as measurement by paxillin staining, was observed near the periphery of the majority of cells on stiff elastic substrates (**Figures 5B, 6**). In contrast, fibroblasts on soft viscoelastic substrates, more reminiscent of normal healthy soft tissue, contained little to no punctate localization of paxillin, which can be attributed to the increase in viscous character (loss modulus) preventing spreading and the formation of larger FAs. Fibroblasts on soft elastic and stiff viscoelastic substrates displayed a mix of punctate paxillin staining and diffuse staining. Cells on Fn9*10 hydrogels, which typically remained rounded regardless of stiffness or viscoelasticity, showed mainly diffuse paxillin staining. Fibroblasts on Fn4G αvβ3-engaging elastic hydrogels also led to a mix of paxillin punctate structures and diffuse staining, similar to those seen with RGD groups. Again, viscoelasticity played a role in suppressing the formation of larger focal adhesions. These findings were also observed quantitatively with fibroblasts on αvβ3-engaging hydrogels (RGD, Fn4G) displaying increased focal adhesion area (**Figure S10**). However, some large, mature FAs were observed for fibroblasts seeded on soft elastic Fn4G hydrogels. Together, these results suggest that preferential αvβ3 binding may facilitate focal adhesion maturation and subsequent actin stress fiber organization and spreading even on soft hydrogels that are more linearly elastic, perhaps mimicking the soft but less viscoelastic mechanical environment observed in active fibroblastic foci in progressive pulmonary fibrosis^25^.

**Figure 6.**
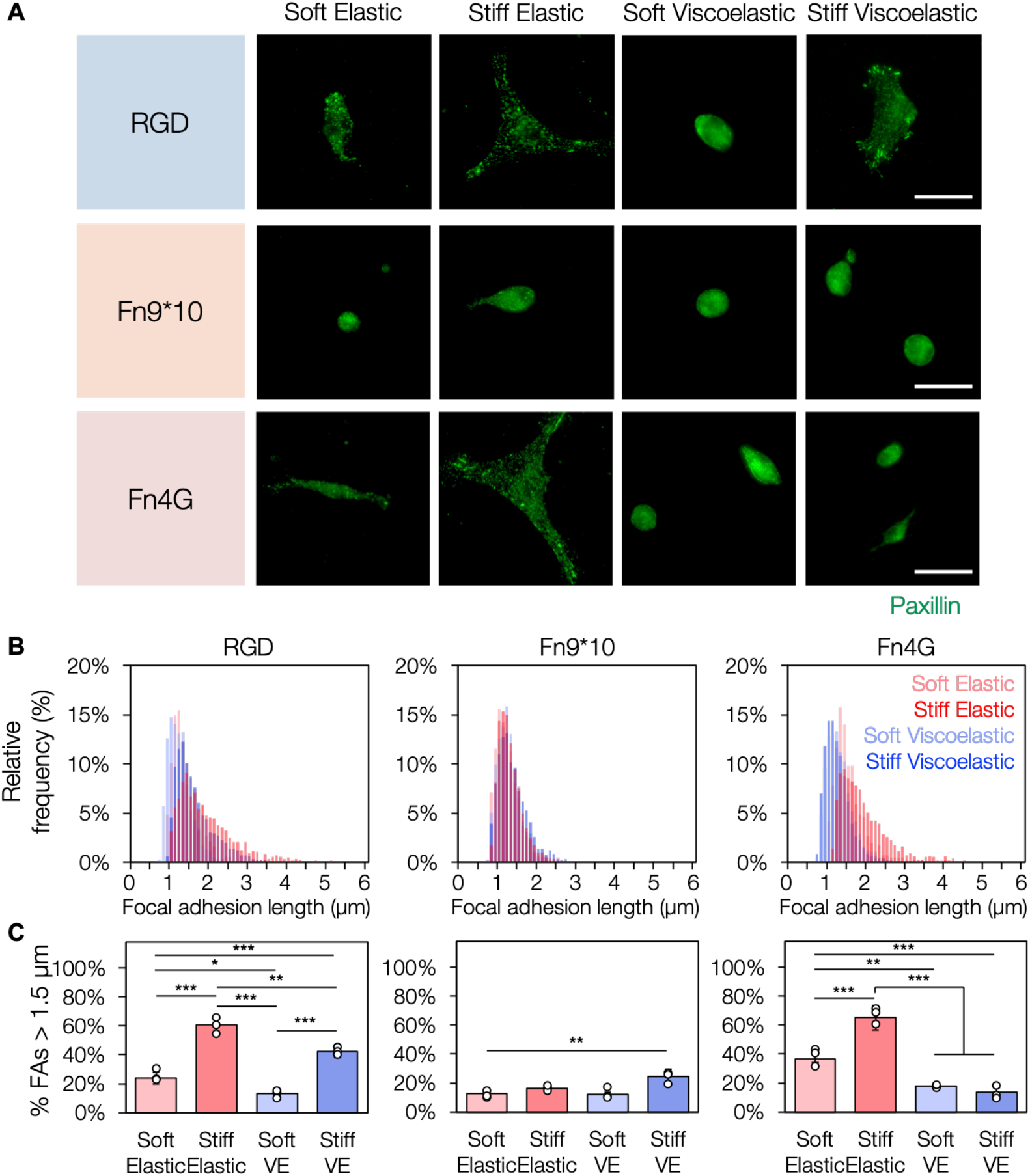
Preferential αvβ3 integrin engagement promotes larger focal adhesion formation. (A) Human lung fibroblasts seeded on hydrogels preferentially binding αvβ3 displayed more punctate paxillin staining on stiff elastic substrates, but viscoelasticity suppressed focal adhesion organization and maturation. *Scale bars:* 50 μm. (B) Histograms of focal adhesion length (determined via quantification of paxillin staining) for fibroblasts cultured on hydrogels for one day. Histograms are grouped by ligand and superimposed to show variance as a function of stiffness and viscoelasticity. (C) The percentages of focal adhesion lengths over 1.5 μm for each hydrogel group. Fibroblasts on Fn9*10-functionalized α5β1-engaging hydrogels had smaller focal adhesions regardless of stiffness and viscoelasticity. *: *P* < 0.05, **: *P* < 0.01, ***: *P* < 0.001; *n* > 180 adhesions from at least 3 hydrogels per experimental group.

## 4. Conclusions

We have described the successful design and implementation of a modular hydrogel platform enabling independent control of covalent crosslinking, incorporation of supramolecular guest-host interactions, and functionalization with cell adhesive groups differentially engaging integrin heterodimers. Hydrogels with stiffnesses approximating normal and fibrotic lung tissue were synthesized in both elastic and viscoelastic forms presenting either RGD or Fn fragments promoting preferential α5β1 or αvβ3 binding. We then showed that fibroblasts seeded on hydrogels preferentially engaging αvβ3 (RGD, Fn4G) generally showed increased spreading, actin stress fiber formation, and focal adhesion size on stiffer elastic hydrogels, but viscoelasticity played a role in suppressing spreading and focal adhesion maturation regardless of adhesive ligand presentation. In particular, fibrosis-associated αv engagement on Fn4G-modified hydrogels promoted increased spread area and focal adhesion size, even on softer elastic materials. Together, these results highlight the importance of understanding the combinatorial role that viscoelastic and adhesive cues play in regulating fibroblast mechanobiology.

## Supporting Information

^1^H NMR spectra for NorHA and CD-HA, MALDI spectra for the adamantane peptide, additional hydrogel mechanical characterization, and additional cell analysis can be found in the Supporting Information.

## Acknowledgments

This work was supported by the UVA Fibrosis Initiative and the NIH (R35GM138187, T32GM008715). The content is solely the responsibility of the authors and does not necessarily represent the official views of the National Institutes of Health.

