## Supplemental Information for "The combined influence of viscoelasticity and adhesive cues on fibroblast spreading and focal adhesion formation"

### Supplementary Figures

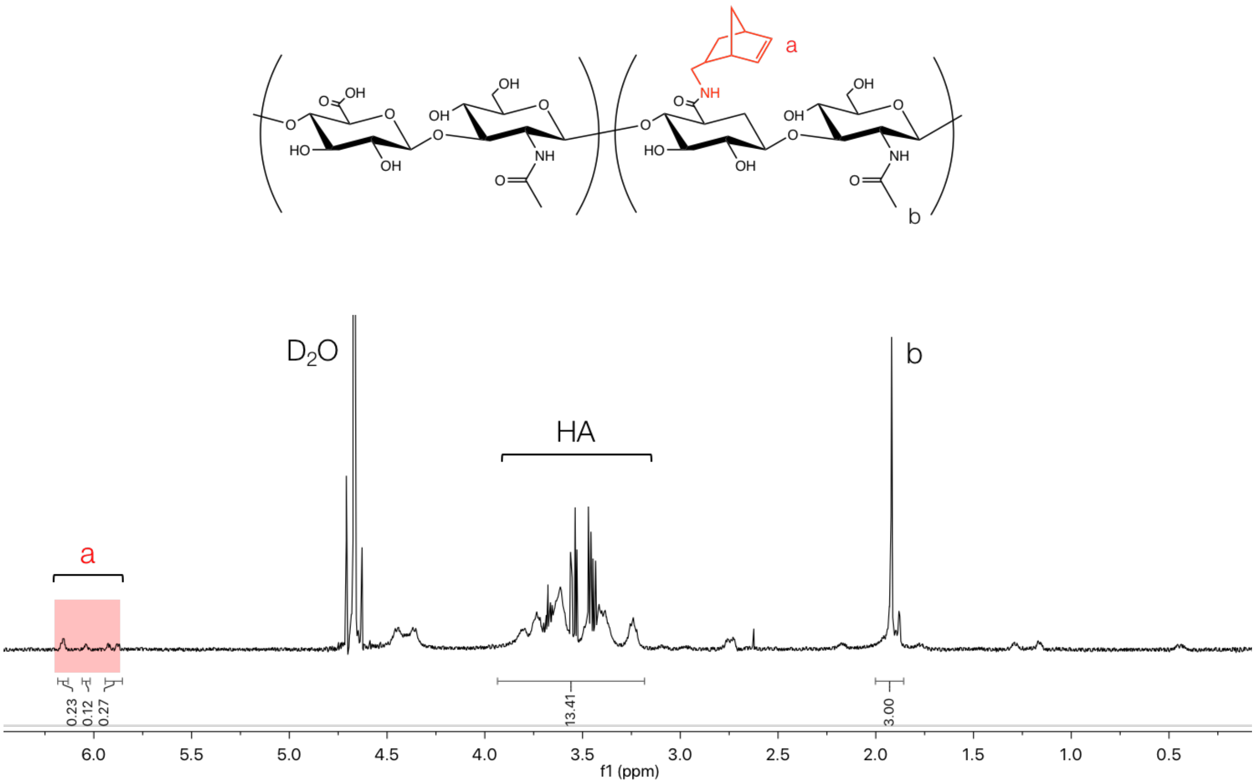


**Figure S1. $^1$H NMR spectrum of norbornene-functionalized hyaluronic acid (NorHA).** The degree of modification, based on norbornene peaks ('a') relative to the methyl peak ('b'), was determined to be 31%.

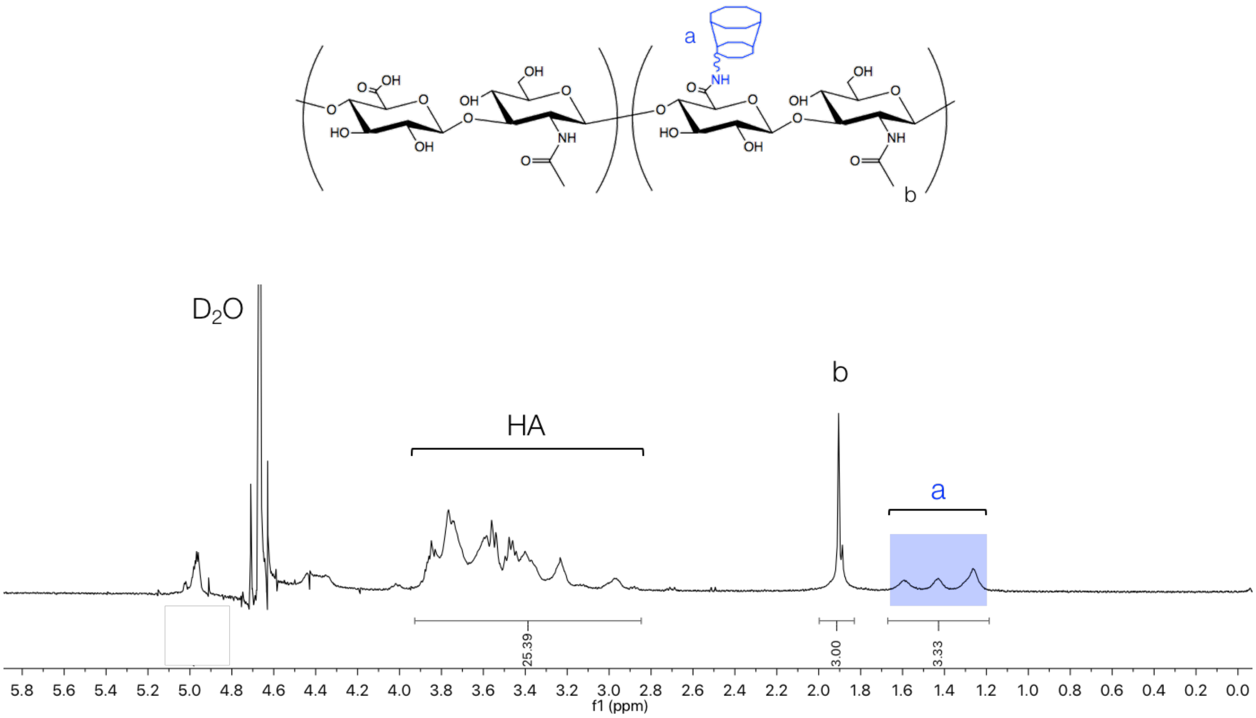


**Figure S2. [1]H NMR spectrum of β-cyclodextrin-functionalized hyaluronic acid (CD-HA).** The degree of modification was determined to be 28%.

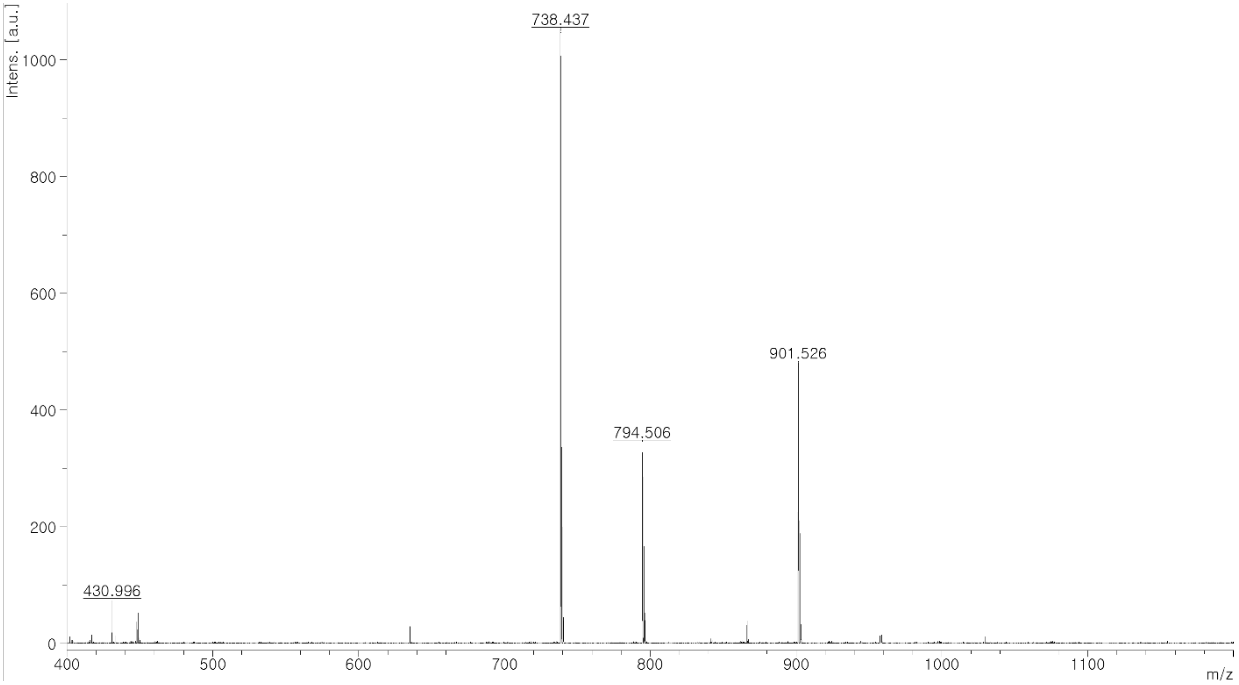


**Figure S3. MALDI spectrum of adamantane (Ad) peptide with the sequence 1-adamantaneacetic acid-KKKCG.** Expected mass: 738.6 g/mol. Actual mass: 738.4 g/mol.

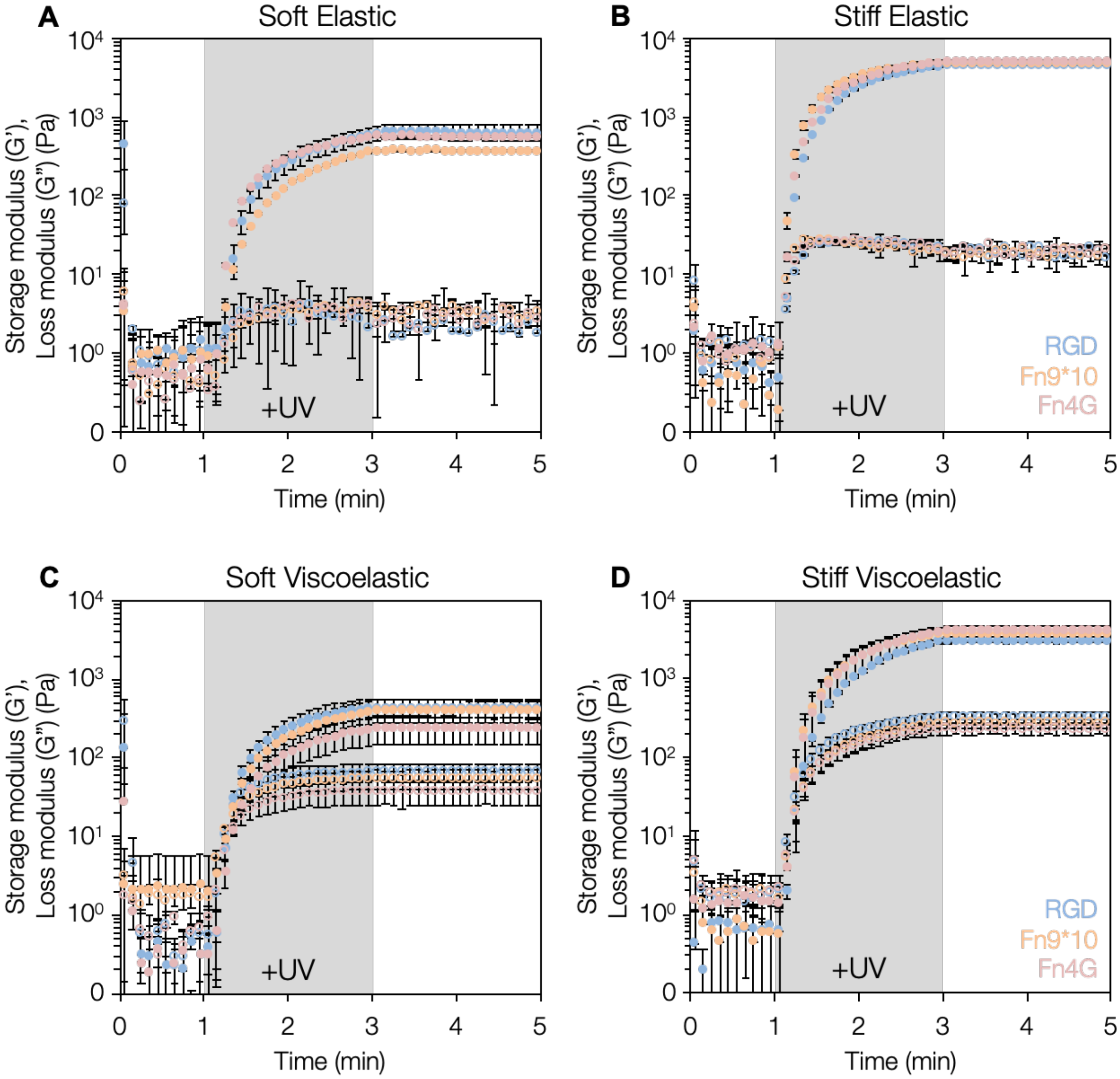


**Figure S4.** ***In situ* gelation of hydrogel groups.** Rheological characterization of elastic and viscoelastic hydrogels representing normal (G' ~ 0.5 kPa, 'soft') and fibrotic (G' ~ 5 kPa, 'stiff') tissue. Viscoelastic groups displayed loss moduli (*G", open circles*) within an order of magnitude of the storage moduli (*G', filled circles*). The gray shaded regions show the 2 minute UV light exposures during gelation. 3 hydrogels were tested per experimental group.

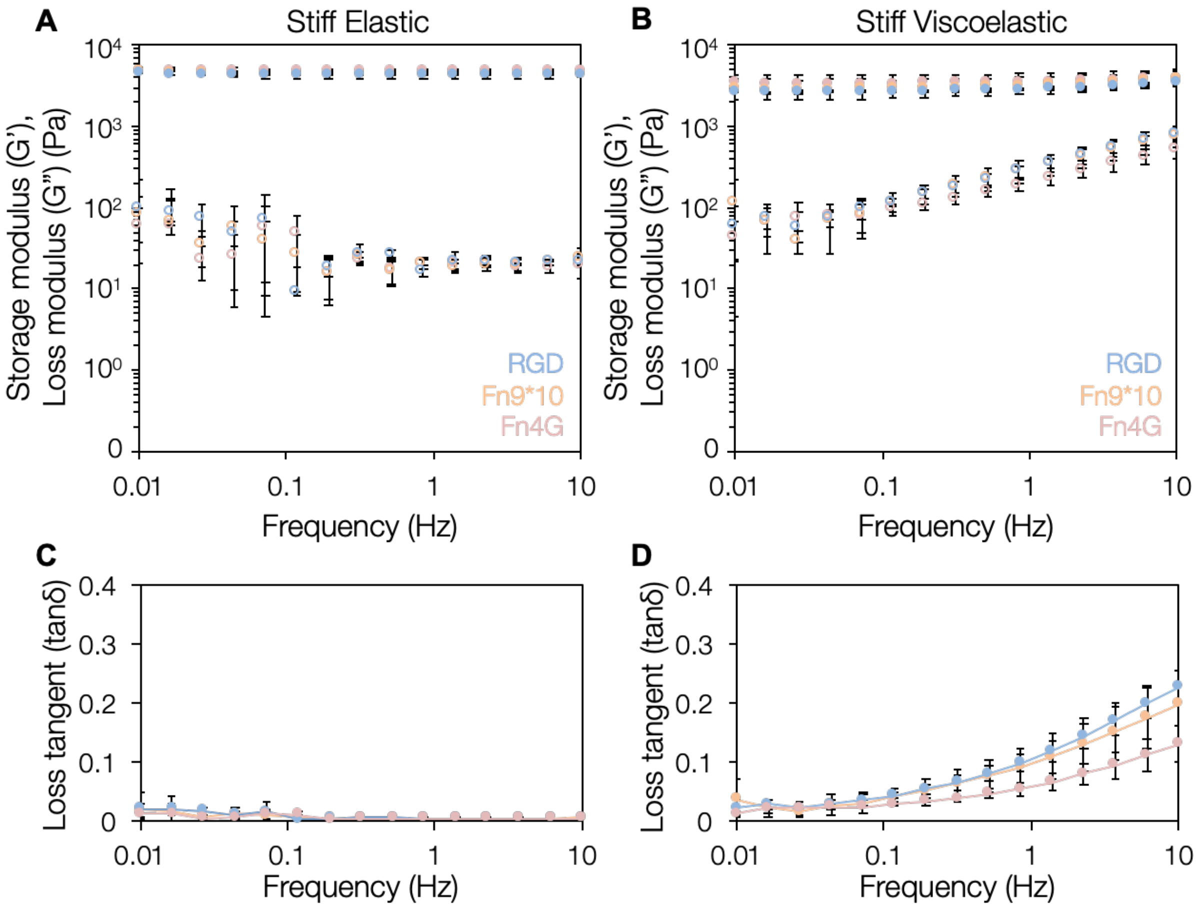


**Figure S5. Rheological behavior of stiff hydrogel groups.** (A) Stiff elastic hydrogels showed frequency-independent behavior with constant loss moduli relative to frequency. (B) Viscoelastic hydrogels displayed frequency-dependent behavior with increasing loss moduli at increasing frequencies. (C) Loss tangent (tanδ, G"/G') values remained relatively constant for all stiff elastic hydrogels. (D) In contrast, loss tangent values were elevated for viscoelastic groups across all frequencies tested and increased at higher frequencies. The soft hydrogel groups can be found in Figure 3.

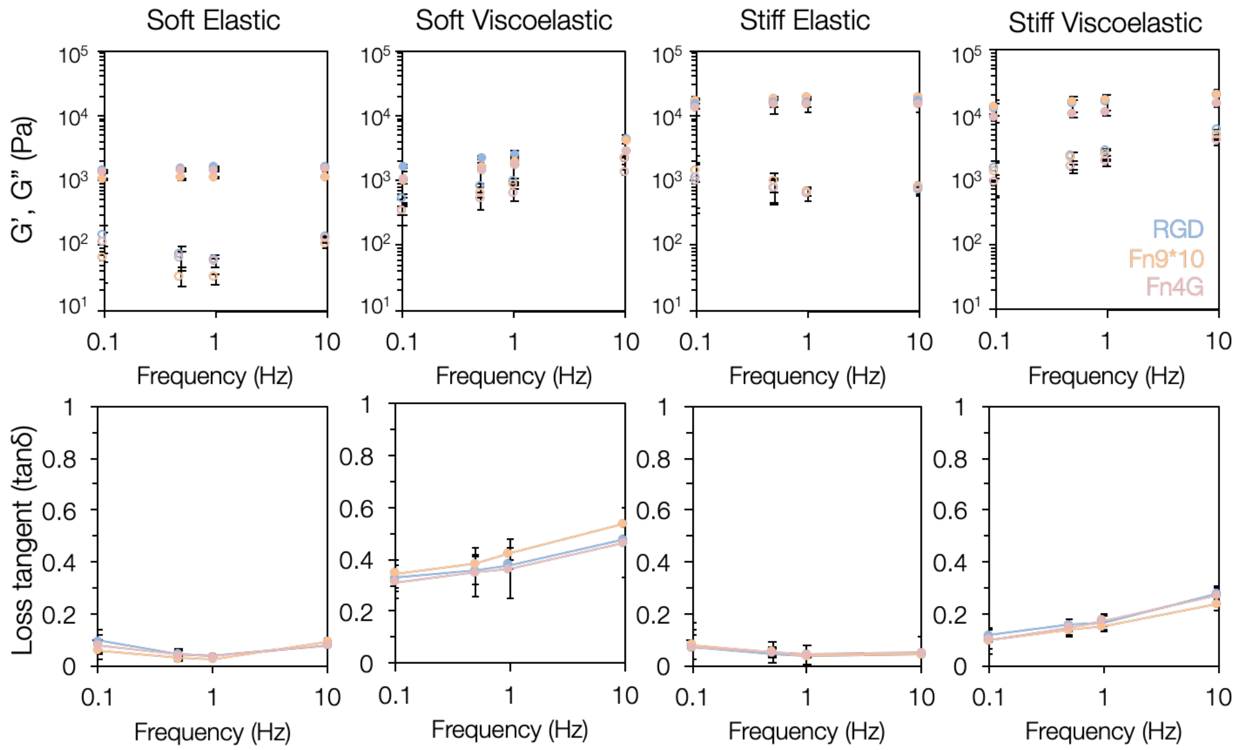


**Figure S6. Mechanical characterization of swollen hydrogels via nanoindentation.** Dynamic mechanical analysis (DMA)-like analysis of PBS-swollen hydrogel groups showed similar frequency-dependent behavior for viscoelastic groups and relatively constant trends for elastic hydrogels.

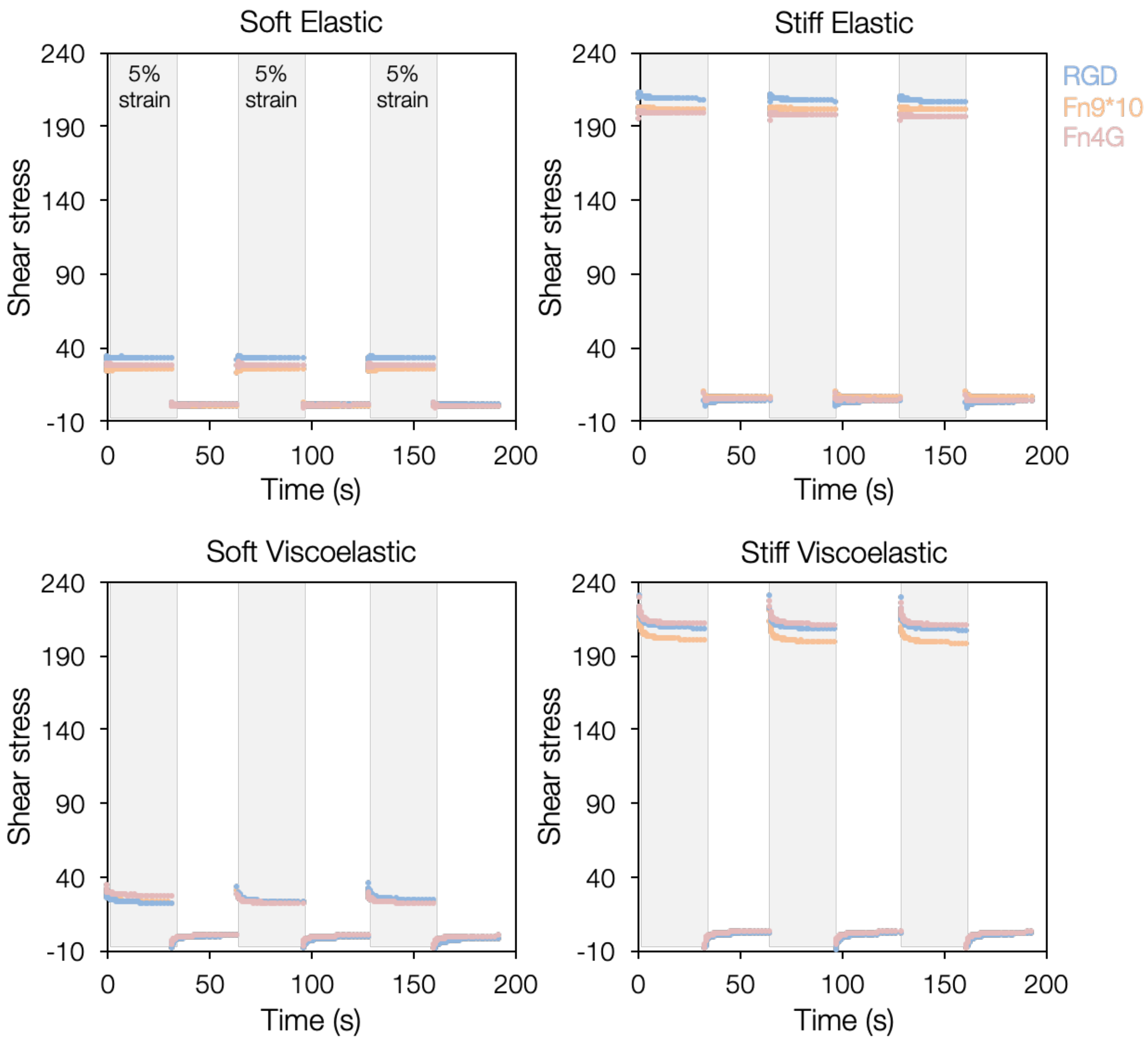


**Figure S7. Stress relaxation and recovery tests.** Cyclic stress relaxation and recovery tests showed full recovery of mechanical properties of hydrogel groups with stress relaxation only occurring in the viscoelastic groups for all ligand types. Strain cycled between 5% (gray bars) and 0.1% (white areas).

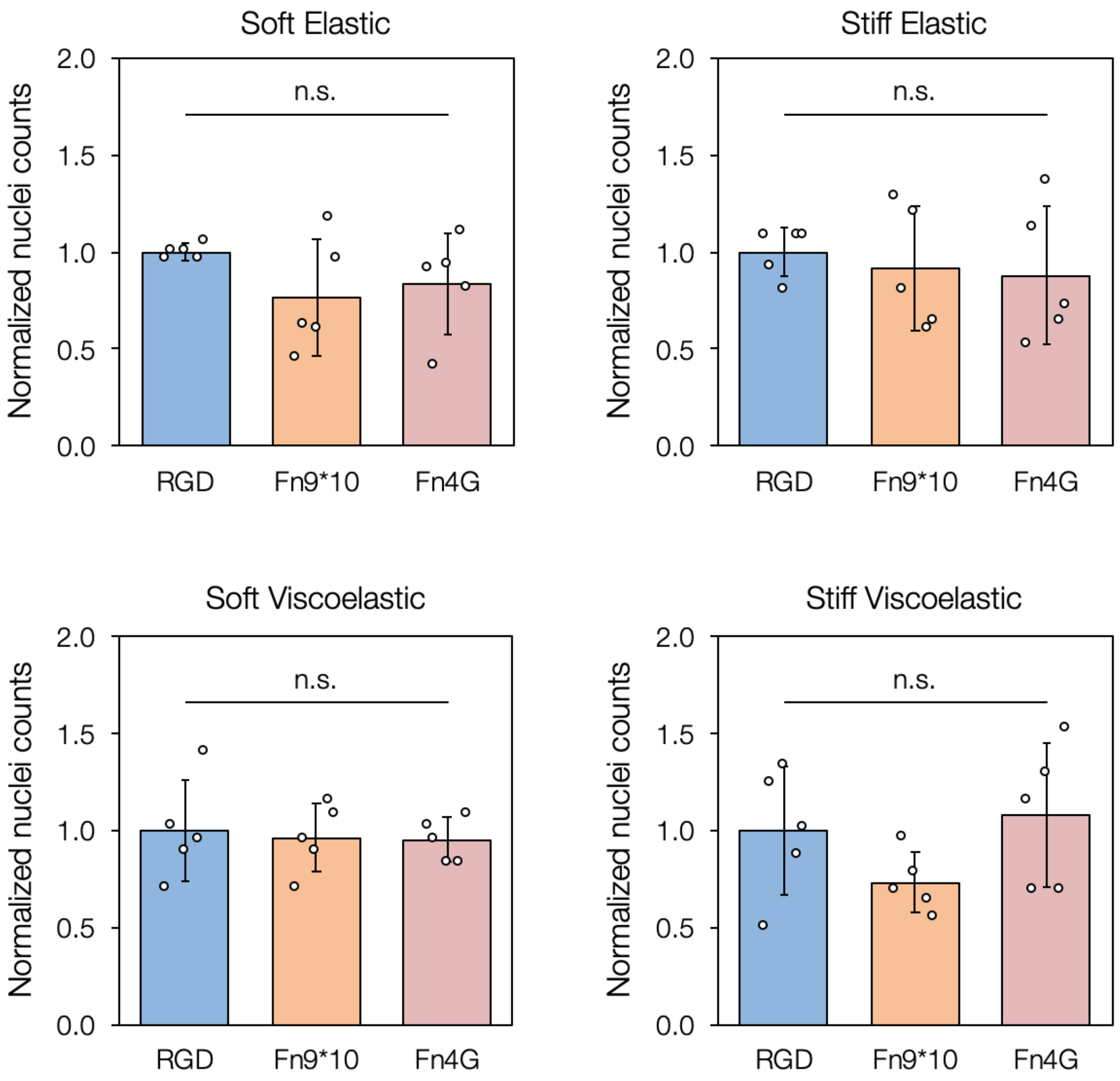


**Figure S8. Fibronectin fragment-functionalized hydrogels support equivalent fibroblast attachment to RGD-modified hydrogels.** Nuclei counts of fibroblasts adhered to all hydrogel experimental groups after one day showed no significant differences between RGD (1 mM) and Fn fragment (2 μM) groups. Nuclei counts were normalized to the RGD groups for each graph. 5 hydrogels were tested per experimental group.

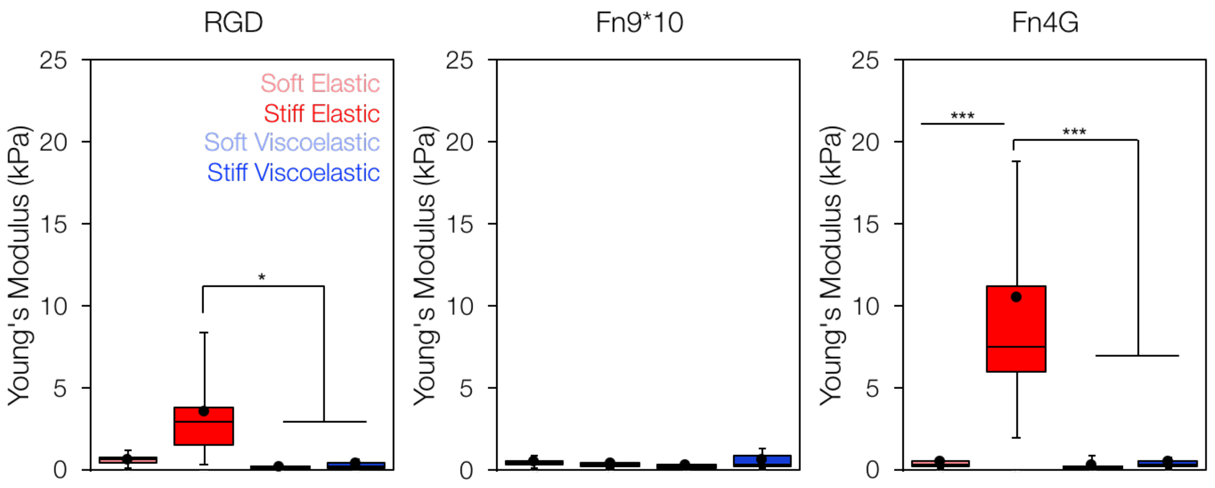


**Figure S9. Nanoindentation measurements of cell stiffness.** Fibroblasts are stiffer on hydrogels promoting αvβ3 engagement on stiff elastic substrates.*: $P < 0.05$, ***: $P < 0.001$; $n$ = 9-19 cells from 3 hydrogels per experimental group.

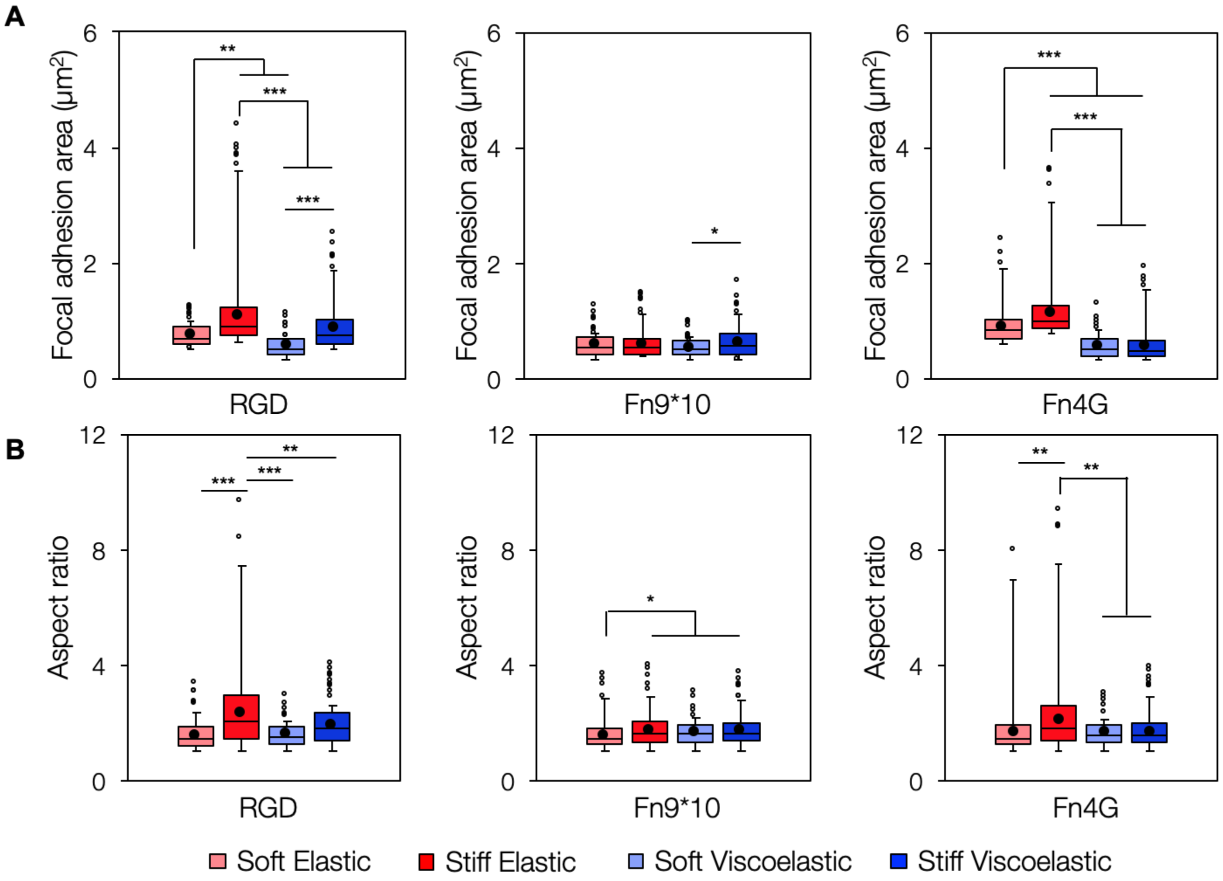


**Figure S10. Focal adhesion area quantification.** A) Human lung fibroblasts on hydrogels preferentially engaging αvβ3 (RGD, Fn4G) displayed increased focal adhesion area as measured by paxillin staining on stiffer, more elastic substrates while fibroblasts on Fn9*10 show reduced focal adhesion size regardless of substrate stiffness or viscoelasticity. B) Focal adhesion aspect ratio quantification showed similar trends to area measurements. Box plots of single cell data show median (*line*), mean (*filled black circle*), and have error bars corresponding to the lower value of either 1.5*interquartile range or the maximum/minimum value, with data points outside the 1.5*interquartile range shown as open circles. *: $P < 0.05$, **: $P < 0.01$, ***: $P < 0.001$; $n >$ 180 adhesions from at least 3 hydrogels per experimental group.
